# A novel regulator of the fungal phosphate starvation response revealed by transcriptional profiling and DNA affinity purification sequencing

**DOI:** 10.1101/2025.04.07.647663

**Authors:** Lori B. Huberman, Vincent W. Wu, David J. Kowbel, Juna Lee, Chris Daum, Vasanth R. Singan, Igor V. Grigoriev, Ronan C. O’Malley, N. Louise Glass

## Abstract

Cells must accurately sense and respond to nutrients to compete for resources and establish growth. Phosphate is a critical nutrient source necessary for signaling, energy metabolism, and synthesis of nucleic acids, phospholipids, and cellular metabolites. During phosphate limitation, fungi import phosphate from the environment and liberate phosphate from phosphate-containing molecules in the cell. In the model filamentous fungus *Neurospora crassa*, the phosphate starvation response is regulated by the conserved transcription factor NUC-1. The activity of NUC-1 is repressed by a complex of the cyclin-dependent kinase MDK-1 and the cyclin PREG when phosphate is plentiful. When phosphate is limiting, NUC-1 repression by MDK-1/PREG is relieved by the cyclin-dependent kinase inhibitor NUC-2. We investigated the global response of *N. crassa* to phosphate starvation. During phosphate starvation, NUC-1 directly activated expression of genes encoding phosphatases, nucleases, and a phosphate transporter and directly repressed genes associated with the ribosome. Additionally, NUC-1 indirectly activated the expression of an uncharacterized transcription factor, which we named *nuc-3*. NUC-3 directly repressed the expression of genes involved in phosphate acquisition and liberation after an extended period of phosphate starvation. Additionally, NUC-3 directly repressed expression of the cyclin-dependent kinase inhibitor *nuc-2*. Thus, through the combination of NUC-3 direct repression of genes in the phosphate starvation response and *nuc-2*, an activator of the phosphate starvation response, NUC-3 serves to act as a brake on the phosphate starvation response after an extended period of phosphate starvation. This braking mechanism could reduce transcription, a phosphate-intensive process, in conditions when phosphate is limiting.

**IMPORTANCE:** Fungi evolved regulatory networks to respond to available nutrients. Phosphate is frequently a limiting nutrient for fungi critical for many cellular functions, including nucleic acid and phospholipid biosynthesis, cell signaling, and energy metabolism. The fungal response to phosphate limitation is important in interactions with plants and animals. We investigated the global transcriptional response to phosphate starvation and the role of a major transcriptional regulator, NUC-1, in the model filamentous fungus *Neurospora crassa*. Our data shows NUC-1 is a bifunctional transcription factor that directly activates phosphate acquisition genes, while directly repressing genes associated with phosphate-intensive processes. NUC-1 indirectly regulates an uncharacterized transcription factor, which we named *nuc-3*. NUC-3 directly represses phosphate acquisition genes and *nuc-2*, an activator of the phosphate starvation response, during extended periods of phosphate starvation. Thus, NUC-3 acts as a brake on the phosphate starvation response to reduce phosphate-intensive activities, like transcriptional activation, when phosphate starvation persists.

## INTRODUCTION

Fungi evolved complex regulatory networks to respond to available nutrients to establish fungal colonies and outcompete other microbes. Phosphate is a critical nutrient for a variety of cellular functions, including nucleic acid, phospholipid, and cellular metabolite biosynthesis, cell signaling, and energy metabolism (1, 2). The ability of fungal cells to respond to phosphate limitation is also important for fungal pathogenesis of plants and animals and establishing symbiotic relationships with plants (3-7).

Fungi have evolved phosphate acquisition pathways to import and metabolize phosphate in phosphate limiting conditions (1, 8, 9). Fungi secrete alkaline phosphatases to liberate inorganic phosphate from the environment (10), while nucleases and acid phosphatases are important in nucleic acid catabolism (2, 11-13). Low affinity phosphate transporters import phosphate into the cell in phosphate-rich environments (14, 15). The expression of high-affinity phosphate transporters is upregulated when environmental phosphate levels are low (16-21). During phosphate abundance, fungi store phosphate as polyphosphate in the vacuole (9, 22, 23). When phosphate is limiting, fungi liberate their vacuolar polyphosphate stores by cleaving polyphosphate and transporting phosphate into the cytoplasm (22-24).

A conserved regulatory pathway activates the expression of genes important for phosphate acquisition during phosphate limitation in fungi (1, 8, 25). Under phosphate-rich conditions, a complex of the cyclin-dependent kinase MDK-1 (NCU07580, previously known as PGOV) in the model filamentous fungus *Neurospora crassa* or Pho85 in the model yeast *Saccharomyces cerevisiae* and the cyclin PREG (NCU01738, *N. crassa*)/Pho80 (*S. cerevisiae*) represses the activity of the basic helix-loop-helix transcription factor, NUC-1 (NCU09315, *N. crassa*)/Pho4 (*S. cerevisiae*), causing NUC-1/Pho4 to localize to the cytoplasm (2, 26-31). When phosphate becomes limiting, the cyclin-dependent kinase inhibitor NUC-2 (NCU11426, *N. crassa*)/Pho81 (*S. cerevisiae*) represses the MDK-1/Pho85 – PREG/Pho80 complex, a process mediated by myo-D-inositol heptakisphosphate (IP_7_) in *S. cerevisiae* and potentially by the MAK-2 (NCU02393) mitogen activated protein kinase in *N. crassa* (32-37). This inhibition enables nuclear localization of NUC-1/Pho4, where it activates phosphate acquisition genes (28, 29).

Although several genes are consistently activated across ascomycete species by NUC-1/Pho4 homologs during phosphate limitation, the extent of the NUC-1/Pho4 regulon differs between species (1, 8). *S. cerevisiae* expresses just over 20 genes during phosphate limitation, while other fungi, including *N. crassa*, have much broader phosphate starvation regulons (32, 38, 39). We set out to interrogate the role of *N. crassa* NUC-1 in the regulation of genes responding to phosphate starvation using a combination of RNA sequencing (RNAseq) and DNA affinity purification sequencing (DAPseq). Our data show that NUC-1 is a bifunctional transcription factor that directly regulates the expression of phosphate transporters, phosphatases, genes involved in nucleic acid, carbohydrate, and lipid metabolism, and genes encoding ribosome-associated proteins or that are involved in ribosome maturation.

We also identified an uncharacterized basic helix-loop-helix transcription factor, which we named NUC-3, that was upregulated during phosphate starvation in a NUC-1-dependent manner. Deletion of *nuc-3* resulted in increased expression of phosphate acquisition genes after 12 hours of phosphate starvation, indicating that NUC-3 represses phosphate acquisition genes once cells have experienced phosphate starvation for an extended time. NUC-3 is conserved in a number of Ascomycete fungi, although no homologs exist in *S. cerevisiae*. NUC-3 directly represses genes involved in phosphate acquisition, liberation of intracellular phosphate stores, lipid metabolism, carbon metabolism, and *nuc-2*, an activator of NUC-1 activity. Our data suggest a mechanism through which fungi place a brake on the phosphate starvation response, repressing gene transcription, a phosphate-intensive cellular process, in phosphate limiting conditions.

## RESULTS

### Genes associated with phosphate acquisition were upregulated and genes associated with the ribosome were downregulated during phosphate starvation

Several previous studies have investigated the global response of *N. crassa* to phosphate limitation (32, 39). However, we hypothesized that additional genes may be regulated during phosphate starvation. Thus, we profiled the transcriptional response of *N. crassa* to phosphate starvation. We grew wild-type cells in media containing 37 mM phosphate for 24h, washed the mycelia, and then exposed cells to either phosphate starvation or 7 mM phosphate for 4h prior to harvesting cells for transcriptional analysis via RNAseq.

The expression of 600 genes was at least 4-fold differentially regulated in wild type cells between phosphate starvation and 7 mM phosphate, with 295 genes downregulated and 305 upregulated in response to phosphate starvation (Fig 1A and Dataset S1). As expected, genes associated with phosphate-intensive cellular activities were downregulated, while genes associated with phosphate acquisition were upregulated. The 295 genes downregulated by at least 4-fold in response to phosphate starvation were enriched for GO categories associated with the ribosome, translation, RNA processing and binding, and branched-chain, lysine, and glutamine metabolic processes (Fig 1B). The majority of downregulated genes were associated with the eukaryotic and mitochondrial ribosomes, rRNA processing, and ribosome assembly – cellular processes that require substantial quantities of phosphate. The 305 genes upregulated by at least 4-fold in response to phosphate starvation were enriched for GO categories associated with phosphate transport, the cell wall, carbon-sulfur lyase activity, cell differentiation, carbohydrate metabolism, and redox reactions (Fig 1C).

**Fig 1.**
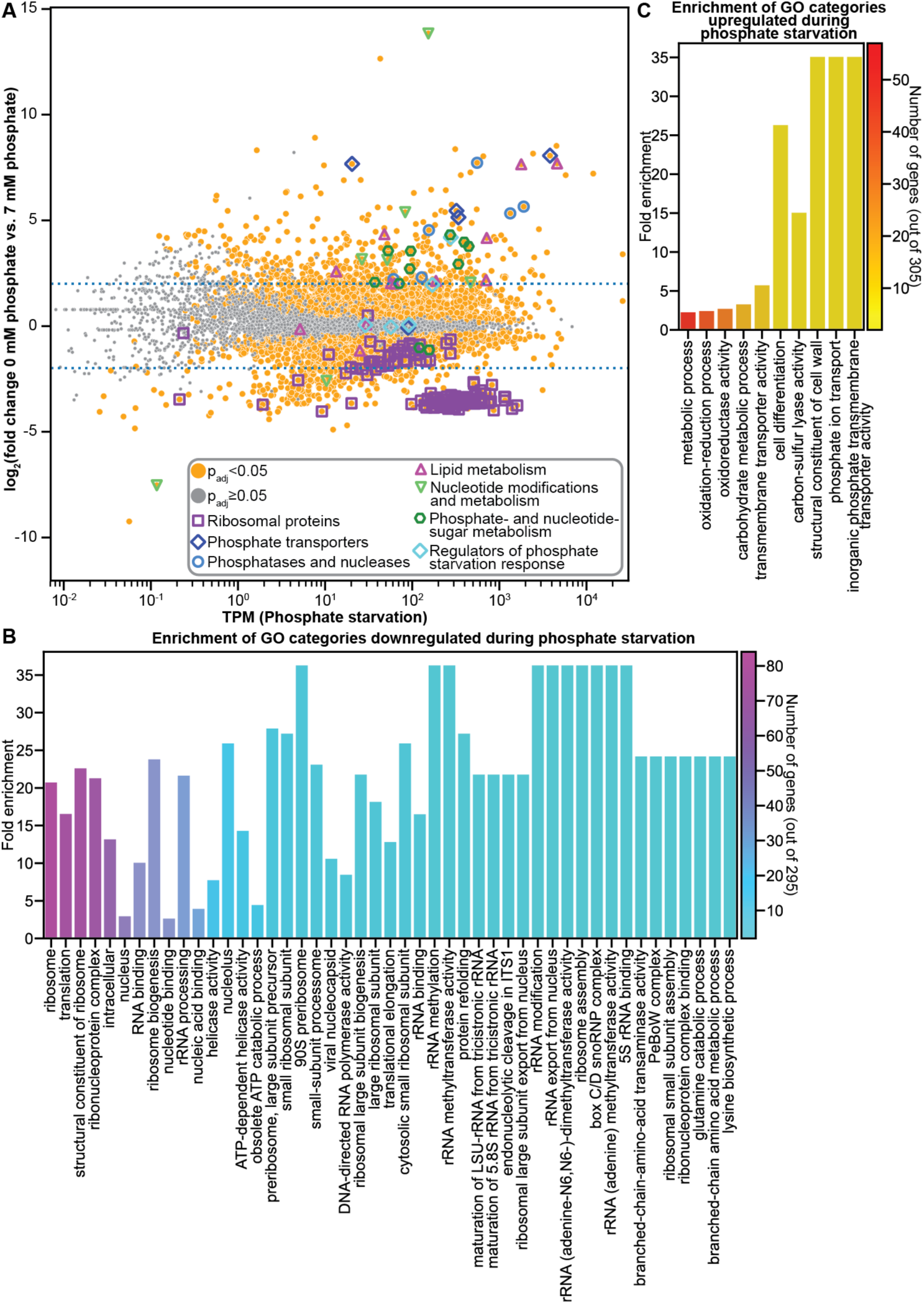
Genes involved in phosphate acquisition and liberation are upregulated and ribosomal proteins are downregulated in response to phosphate starvation. **(A)** Differential expression analysis of wild type cells exposed to phosphate starvation as compared to 7 mM phosphate. Genes with significant differential expression (p_adj_ < 0.05) are indicated with orange circles. Genes without a significant difference in expression (p_adj_ ≥ 0.05) are indicated in grey. Genes encoding predicted ribosomal proteins are indicated with purple squares. Genes encoding known or predicted phosphate transporters are indicated with dark blue diamonds. Genes encoding known or predicted phosphatases and nucleases are indicated with light blue circles. Genes predicted to be involved in lipid metabolism are indicated with magenta triangles. Genes predicted to play a role in nucleotide modification or metabolism are indicated with light green triangles. Genes involved in phosphate-sugar and nucleotide-sugar metabolism are indicated with dark green hexagons. Genes encoding regulators of the phosphate starvation response are indicated with light blue diamonds. Dotted blue lines indicated a 4-fold change in expression. **(B)** Fold enrichment of genes significantly downregulated by at least 4-fold during phosphate starvation as compared to 7 mM phosphate in significantly enriched (p_adj_ < 0.05) gene ontology (GO) categories. The number of genes significantly downregulated by at least 4-fold found in each category is indicated by the color of the bar. **(C)** Fold enrichment of genes significantly upregulated by at least 4-fold during phosphate starvation as compared to 7 mM phosphate in significantly enriched (p_adj_ < 0.05) GO categories. The number of genes significantly upregulated by at least 4-fold found in each category is indicated by the color of the bar. GO enrichment analysis was calculated using FungiFun 2.2.8 (https://elbe.hki-jena.de/fungifun/).

The most highly upregulated genes included genes involved in phosphate acquisition. The alkaline phosphatase *pho-2* (NCU01376) was upregulated by 368-fold, while the acid phosphatase *pho-3* (NCU08643) was upregulated by 286-fold. Several additional genes may also act to acquire phosphate from the environment through phosphatase or nuclease activity. For example, NCU09631 contains an alkaline phosphatase domain and was upregulated by 211-fold. Several uncharacterized nucleases were also upregulated by at least 20-fold, including an extracellular deoxyribonuclease (NCU09525), the ribonuclease T1 *grn* (NCU01045), and the nuclease *nuc-14* (NCU09788). We also saw upregulation of the predicted histidine pyrophosphatase *pht-1* (NCU06351) and the inorganic phosphatase *ipp-2* (NCU08703) (Fig 1A and Dataset S1).

Transporters are required to import phosphate into the cell. The high-affinity phosphate transporters *pho-4* (NCU09564) and *pho-5* (NCU08325) were upregulated by 204-fold and 267-fold, respectively, under phosphate starvation. Several additional transporters that may play a role in phosphate transport were also upregulated (Fig 1A and Dataset S1). The chromate ion transporter *trm-50* (NCU01055), which may also be able to transport phosphate, was upregulated by 44-fold. Aside from the Major Facilitator Superfamily (MFS) transporter *pho-5*, 13 additional MFS transporters were upregulated in response to phosphate starvation. These 13 MFS transporters included NCU09767, which has homology to phosphate transporters, the predicted phospholipid transporter *mfs-10* (NCU04809), two transporters with homology to nicotinic acid transporters (NCU09027 and NCU08715), and a predicted pantothenate transporter (*mfs-16* [NCU00782]) (Dataset S1). The expression of carbohydrate transporters was also upregulated, including: high-affinity glucose transporters [*sut-9* (NCU04963) and *hgt-1* (NCU10021)], the L-arabinose transporter *lat-1* (NCU02188), the rhamnose transporter *sut-28* (NCU05897) (40), and *sut-18* (NCU09287). Three additional MFS transporters were also upregulated: NCU04446, *asd-3* (NCU05597), and *mfs-24* (NCU07343). Other upregulated transporters included the ATP-binding cassette (ABC) superfamily transporter *mig-12* (NCU09830), the P-type ATPase *ph7* (NCU08147), the ion-translocating microbial rhodopsin *nop-1* (NCU10055), and the iron/lead transporter NCU09210 (Dataset S1).

During phosphate starvation, fungi not only import phosphate from the environment, they also liberate phosphate from phosphate-containing molecules. The 3’(2’),5’-bisphosphate nucleotidase *inl-13* (NCU04069), which releases phosphate from adenosine 3’,5’-bisphosphate was the most upregulated gene during phosphate starvation with a 14,000-fold increase in expression (Fig 1A and Dataset S1). Other genes whose expression was upregulated by at least 4-fold during phosphate starvation may also play a role in releasing phosphate from nucleotides, including NCU05005 and NCU09856, which may have ATP hydrolysis activity, a potential NADH pyrophosphatase (NCU01127), and the predicted uracil phosphoribosyl transferase *uc-8* (NCU06261) (Fig 1A and Dataset S1).

During phosphate starvation, fungi release phosphate from phospholipids and replace phospholipids with betaine lipids (41). A glycerophosphoryl diester phosphodiesterase, which cleaves phosphate from phospholipids, *gdp-1* (NCU10038), and the betaine lipid synthase *bet-6* (NCU03032) were both upregulated by more than 200-fold during phosphate starvation (Fig 1A and Dataset S1). Several other genes whose expression was upregulated may also play a role in harvesting phosphate from phospholipids, including three genes that may have lipase activity: NCU05859, *cea-6* (NCU04930), and NCU09526. The expression of genes with other potential lipid-associated roles were also upregulated, including the predicted phosphatidylethanolamine binding protein NCU01112; *chol-11* (NCU02302), which may play a role in phosphatidylethanolamine synthesis; and the glycerophosphocholine phosphodiesterase *gde-1* (NCU01747), which hydrolyzes glycerophosphocholine to choline and glycerol phosphate for use as a phosphate source and is homologous to *S. cerevisiae GDE1*, a gene activated in response to phosphate (38, 42) (Fig 1A and Dataset S1).

Phosphate is also present in metabolic intermediates of carbohydrate utilization. We identified upregulated genes with predicted roles in modifying nucleotide sugars and phosphorylated sugars, suggesting that this may be another method through which phosphate is liberated during phosphate starvation. Three genes whose expression was upregulated by at least 6-fold may play roles in nucleotide sugar metabolism: the predicted UPD-galactose-4-epimerase *gae-1* (NCU08549); NCU09906, which may play a role in transferring sugars from UPD-sugars; and *nmr-3* (NCU09403), which has homology to nucleoside diphosphate-sugar 4 epimerases (Fig 1A and Dataset S1). The expression of the mannose-6-phosphate isomerase *man-2* (NCU02322), the fructose bisphosphatase *fbp-1* (NCU04797), the cellobionic acid phosphorylase *ndvB* (NCU09425), the potential fructose-1,6-phosphate aldolase *tqaM* (NCU01072), and two predicted phosphoketolases (*phk-1* [NCU05151] and *phk-2* [NCU06123]) were also upregulated by at least 4-fold in response to phosphate starvation (Fig 1A and Dataset S1).

### Many of the genes involved in the response to phosphate starvation were regulated in a NUC-1-dependent fashion

In *N. crassa*, the transcription factor NUC-1 is required for the response to phosphate starvation (27). However, a global investigation of the role of NUC-1 in gene regulation has not been undertaken. To address this question, we inoculated wild type (*nuc-1^+^*) and Δ*nuc-1* cells in 37 mM phosphate and then shifted the mycelial mass to media lacking phosphate for 4h prior to harvesting the cells for RNAseq. The expression of 162 genes was differentially expressed by at least 4-fold in the Δ*nuc-1* mutant as compared to *nuc-1^+^* cells during phosphate starvation, with 60 genes downregulated and 102 genes upregulated in Δ*nuc-1* cells as compared to *nuc-1^+^* cells. (Fig 2A and 2B). The 102 genes upregulated in Δ*nuc-1* cells as compared to wild type cells in response to phosphate starvation were enriched for GO categories associated with ribosome biogenesis and assembly, rRNA transcription and processing, ribonucleoprotein complex, the nucleolus, and folate synthesis (Fig S1A). Of the 102 genes upregulated in cells lacking *nuc-1* as compared to wild type cells, 94 also met our stringent threshold for 4-fold downregulation in wild type during phosphate starvation as compared to 7 mM phosphate (Fig 2A). The vast majority of these 102 genes are associated with the ribosome, ribosome biogenesis, or transcription, phosphate-hungry cellular activities (Dataset S1).

**Fig 2.**
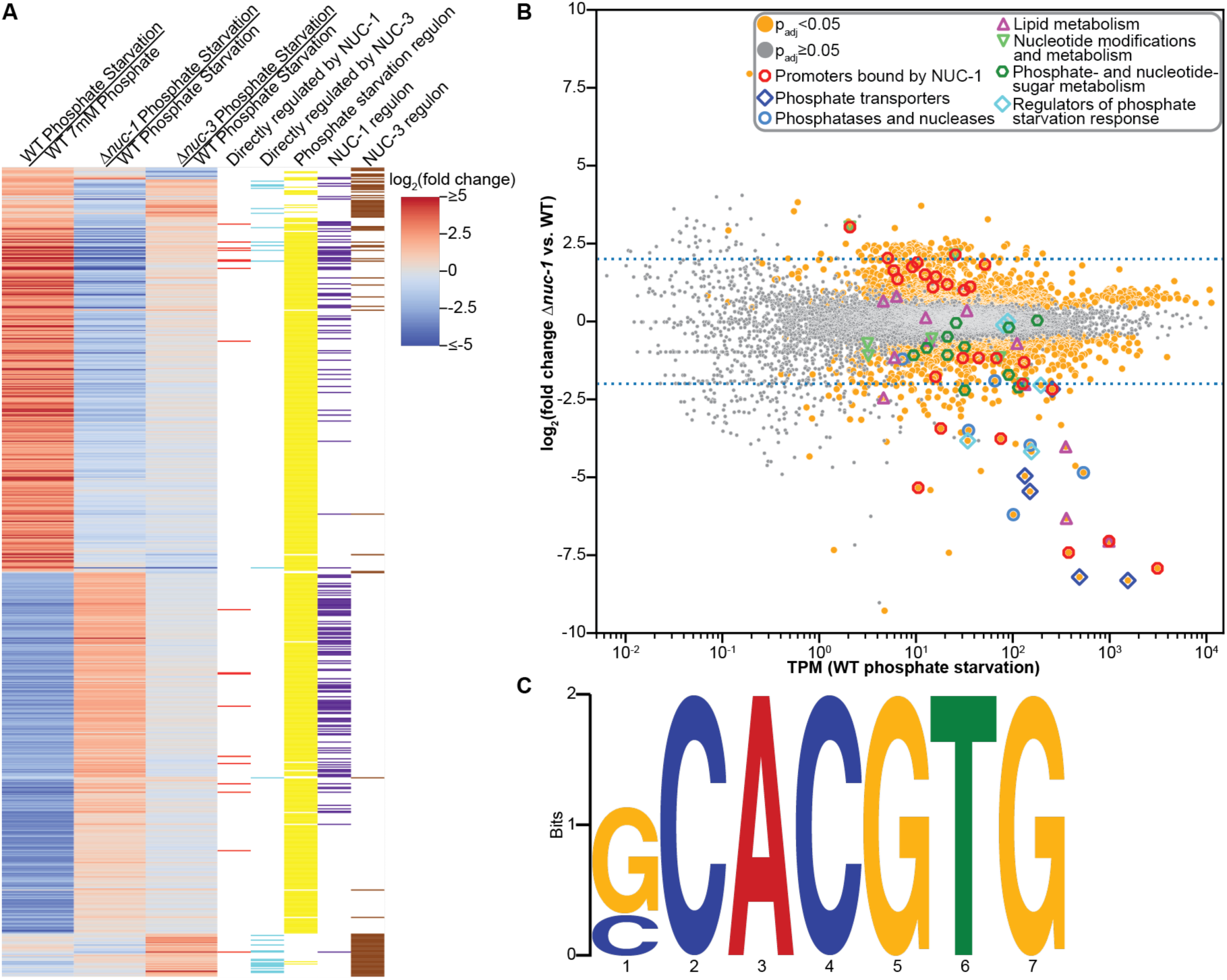
The transcription factor NUC-1 directly activates genes necessary for phosphate acquisition and directly represses genes associated with the ribosome in response to phosphate starvation. **(A)** Heatmap showing the log_2_(fold change in expression) of the indicated comparisons of all genes differentially expressed by at least 4-fold in wild type (WT) cells exposed to phosphate starvation as compared to 7 mM phosphate for 4 h, differentially expressed by at least 4-fold in the Δ*nuc-1* mutant as compared to wild type (*nuc-1^+^*) cells exposed to phosphate starvation for 4 h, or differentially expressed by at least 2-fold in the Δ*nuc-3* mutant as compared to wild type (*nuc-3^+^*) cells exposed to phosphate starvation for 12 h. Red bars indicate genes whose promoter was bound by NUC-1 and were differentially expressed by at least 2-fold in the Δ*nuc-1* mutant as compared to wild type (*nuc-1^+^*) cells exposed to phosphate starvation for 4 h. Cyan bars indicate genes whose promoter was bound by NUC-3 and were differentially expressed by at least 2-fold in the Δ*nuc-3* mutant as compared to wild type (*nuc-3^+^*) cells exposed to phosphate starvation for 12 h. Yellow bars indicate genes that were differentially expressed by at least 4-fold in wild type cells exposed to phosphate starvation as compared to 7 mM phosphate for 4 h. Purple bars indicate genes that were differentially expressed by at least 4-fold in the Δ*nuc-1* mutant as compared to wild type (*nuc-1^+^*) cells exposed to phosphate starvation for 4 h. Brown bars indicate genes that were differentially expressed by at least 2-fold in the Δ*nuc-3* mutant as compared to wild type (*nuc-3^+^*) cells exposed to phosphate starvation for 12 h. **(B)** Differential expression analysis of Δ*nuc-1* cells as compared to wild type (*nuc-1^+^*) cells exposed to phosphate starvation. Genes with significant differential expression (p_adj_ < 0.05) are indicated with orange circles. Genes without a significant difference in expression (p_adj_ ≥ 0.05) are indicated in grey. Genes whose promoters were bound by NUC-1 and were differentially expressed by at least 2-fold in the Δ*nuc-1* mutant as compared to wild type (*nuc-1^+^*) cells exposed to phosphate starvation are indicated with red octagons. Genes encoding known or predicted phosphate transporters are indicated with dark blue diamonds. Genes encoding known or predicted phosphatases and nucleases are indicated with light blue circles. Genes predicted to be involved in lipid metabolism are indicated with magenta triangles. Genes predicted to play a role in nucleotide modification or metabolism are indicated with light green triangles. Genes involved in phosphate-sugar and nucleotide-sugar metabolism are indicated with dark green hexagons. Genes encoding regulators of the phosphate starvation response are indicated with light blue diamonds. Dotted blue lines indicated a 4-fold change in expression. Two genes expressed at transcripts per million (TPM) below 10 whose expression was not significantly different in Δ*nuc-1* cells as compared to wild type (*nuc-1^+^*) cells exposed to phosphate starvation but had a log_2_(fold change in expression) of more than 20-fold (NCU00695 and NCU09638) were left off the scatterplot to better visualize the expression of the other genes. The expression and fold change of these genes can be seen in Dataset S1. **(C)** NUC-1 consensus DNA binding motif (E-value = 2.2 x 10^-9^) of NUC-1 promoter binding sites in genes differentially expressed by at least 2-fold in Δ*nuc-1* cells as compared to wild type cells during phosphate starvation built using MEME version 5.5.5 (78).

The 60 genes downregulated in Δ*nuc-1* as compared to wild type cells in response to phosphate starvation were enriched for GO categories associated with phosphate and chromate transport, nuclease and phosphatase activity, phosphoric diester hydrolase activity, oxidation-reduction processes, and formate dehydrogenase activity (Fig S1B). Most genes (53 of 60 genes) downregulated in Δ*nuc-1* relative to *nuc-1^+^* cells in response to phosphate starvation by at least 4-fold also met our stringent threshold for genes activated in response to phosphate starvation in wild type cells (Fig 2A). Two of the remaining 7 genes included the cyclin-dependent kinase inhibitor *nuc-2*, critical for activation of the phosphate starvation response in *N. crassa* (33) and the MFS phosphate transporter *pho-7* (NCU07375) (Dataset S1).

The 53 genes whose expression was both downregulated in Δ*nuc-1* cells as compared to *nuc-1^+^* cells in response to phosphate starvation and upregulated in response to phosphate starvation as compared to 7 mM phosphate in wild type cells included a number of genes known to play a role in phosphate acquisition: *pho-2*, *pho-3*, *pho-4*, *pho-5*, and *gdp-1* (Fig 2B and Dataset S1). Several predicted nucleases that were activated in response to phosphate starvation in wild type cells were downregulated in Δ*nuc-1* cells relative to *nuc-1^+^*cells, including NCU09631, NCU09525, *grn*, and *nuc-14*. The transporters *trm-50*, NCU09767, *mfs-10*, *sut-28*, and *ph7* were also upregulated in wild type cells in a NUC-1-dependent fashion during phosphate starvation. Additionally, the expression of the predicted uracil phosphoribosyl transferase *uc-8*, which may play a role in releasing phosphate from nucleotides, as well as the predicted UPD-galactose-4-epimerase *gae-1* and the fructose bisphosphatase *fbp-1*, was downregulated in Δ*nuc-1* cells as compared to *nuc-1^+^* cells in response to phosphate starvation. The release of phosphate from phospholipids and activation of betaine lipid synthesis appeared to be repressed in Δ*nuc-1* cells relative to *nuc-1^+^* cells during phosphate starvation, given the downregulation of *bet-6* and *gde-1* (Fig 2B and Dataset S1). Overall, comparing the transcriptional profiling of wild type cells exposed to phosphate starvation as compared to 7 mM phosphate and wild type as compared to Δ*nuc-1* cells during exposure to phosphate starvation demonstrated that while NUC-1 regulates many of the genes involved in the phosphate starvation response, there may also be other transcriptional regulators as the phosphate starvation regulon in wild type cells involved substantially more genes than were in the NUC-1 regulon (Fig 2A).

### NUC-1 directly activated genes involved in phosphate acquisition and liberation and directly repressed genes associated with phosphate-intensive cellular processes

Genes differentially expressed in Δ*nuc-1* cells as compared to *nuc-1^+^* cells during phosphate starvation could be due to direct or indirect regulation by NUC-1. To distinguish between these two possibilities, we measured NUC-1 promoter binding using DAPseq, an *in vitro* method to identify NUC-1 DNA binding sites (40, 43-45). We identified 380 genes with 281 NUC-1 binding sites within 3,000 bp upstream of their translational start site (Dataset S2). Since DAPseq is an *in vitro* technique to measure DNA binding, it is possible that not all genes with NUC-1 promoter binding sites identified via DAPseq were regulated by NUC-1 *in vivo*. Thus, we filtered our DAPseq data for genes that were also at least 4-fold differentially expressed between *nuc-1^+^*cells and Δ*nuc-1* cells during phosphate starvation.

Eleven genes were both at least 4-fold differentially expressed between *nuc-1^+^* cells and Δ*nuc-1* cells and had promoters bound by NUC-1 *in vitro*. Of these, eight genes were activated by NUC-1 and three were repressed by NUC-1 in response to phosphate starvation (Fig 2A, 2B, and Dataset S1). Given this extremely limited set of direct NUC-1 targets, we dropped our threshold for NUC-1-regulation to at least a 2-fold change in the RNAseq comparison between *nuc-1^+^* cells and Δ*nuc-1* cells exposed to phosphate starvation. This reduced threshold increased the number of genes bound and regulated by NUC-1 to 27, with 13 directly activated by NUC-1 and 14 directly repressed by NUC-1 (Fig 2B and Dataset S1).

Four genes bound and directly activated by NUC-1 have a clear connection to phosphate acquisition: *pho-2*, which was previously shown to have a NUC-1 promoter binding site (46); *pho-3* (13); *gdp-1*; and *pho-7* (Fig 2B and Dataset S1). NUC-1 also bound and directly activated *uc-8* and NCU03546, a predicted polyphosphate kinase. Homologs of NCU03546 are important in synthesis of IP_7_, which increases during phosphate starvation and is involved in regulation of the *S. cerevisiae* Pho4p protein (34, 47). The roles of the other seven genes bound and directly activated by NUC-1 during the phosphate starvation response were somewhat less clear. These genes included the metallo-β-lactamase superfamily protein *mbl-5* (NCU07133); a LysM domain containing protein (NCU07486); an F-box domain-containing protein (NCU05581); L-galactonate dehydratase (NCU07064); a NipSnap family protein (NCU08092); the 3-ketoacyl-acyl carrier protein reductase *cel-3* (NCU09473); and a hypothetical protein (NCU07485) (Fig 2B and Dataset S1). To our surprise, although a prior study found two NUC-1 binding sites in the promoter of the phosphate transporter *pho-4* by measuring *in vitro* binding of a truncated NUC-1 protein, we did not identify NUC-1 bound to the *pho-4* promoter using DAPseq with full-length NUC-1 (46).

Most of the 14 genes bound and repressed by NUC-1 were involved in phosphate-intensive cellular processes. Nine were associated with the ribosome or ribosome biogenesis: a predicted pre-rRNA processing protein (NCU00059); the RNA-3’-phosphate cyclase *rbg-27* (NCU02675), whose homologs play roles in rRNA processing (48); the 60S ribosomal protein L20 *mrp-29* (NCU04225); the predicted ribosomal protein eL24 (NCU02644); the 20S pre-rRNA D-site endonuclease NOB1 *rbg-39* (NCU08904); the predicted ribosome biogenesis protein Alb1 (NCU01458); a predicted 25S rRNA adenine-N_1_ methyltransferase (NCU06227); the predicted DNA-directed RNA polymerase I subunit RPA43 (NCU07166); and a predicted RNA polymerase I specific transcription initiation factor RRN3 superfamily member (NCU09247) (Fig 2B and Dataset S1). Three additional genes may play a role in transcription or RNA metabolism, including: the deoxyribose-phosphate aldolase *dpa-2* (NCU00597); a pentatricopeptide repeat protein NCU02674 that is homologous to a gene that may play a role in mRNA metabolism in mammals (49, 50); and NCU07419, an F-box domain containing protein that is homologous to histone demethylases involved in transcriptional activation (51). The GDSL lipase *lip-8* (NCU06364) and the methionine permease *aap-7* (NCU04942) were also directly repressed by NUC-1 (Fig 2B and Dataset S1). We used the NUC-1 promoter binding sites in these 27 genes to identify the consensus binding motif GCACGTG (e-value = 2.2×10^-9^) (Fig 2C). This NUC-1 consensus binding motif is extremely similar to the previously predicted NUC-1 binding motif, CACGTG (46).

### The uncharacterized transcription factor NCU03077 repressed expression of the alkaline phosphatase pho-2 during phosphate starvation

A single transcription factor, NCU03077, was downregulated by at least 4-fold in Δ*nuc-1* cells relative to *nuc-1^+^* cells during phosphate starvation and upregulated in wild type cells exposed to phosphate starvation as compared to 7 mM phosphate (Fig 1A, 2B, and Dataset S1). NCU03077 is uncharacterized, and, like NUC-1, encodes a basic helix-loop-helix transcription factor. Homologs of NCU03077 exist within the Sordariomycetes and other Pezizomycete species but are not present in *S. cerevisiae* (Fig S2).

We hypothesized that NCU03077 might play a role in regulating the expression of phosphate utilization genes. To test this hypothesis, we measured the expression of the alkaline phosphatase *pho-2* in wild type and ΔNCU03077 cells from 2h-24h after shifting cells from 37 mM phosphate to phosphate starvation. In wild type cells, the expression of *pho-2* increased rapidly after a shift to phosphate starvation, peaking at 8h post-shift to phosphate starvation and then dropping to a steady-state expression level, which was maintained from 10h-24h post-shift to phosphate starvation (Fig 3A). The expression of *pho-2* in cells lacking NCU03077 was similar to wild type cells in the first 4h after a shift to phosphate starvation but reached a higher 8h peak and was maintained at a significantly higher level of expression than in wild type cells (Fig 3A). These data suggested that NCU03077 was required to repress expression of *pho-2*.

**Fig 3.**
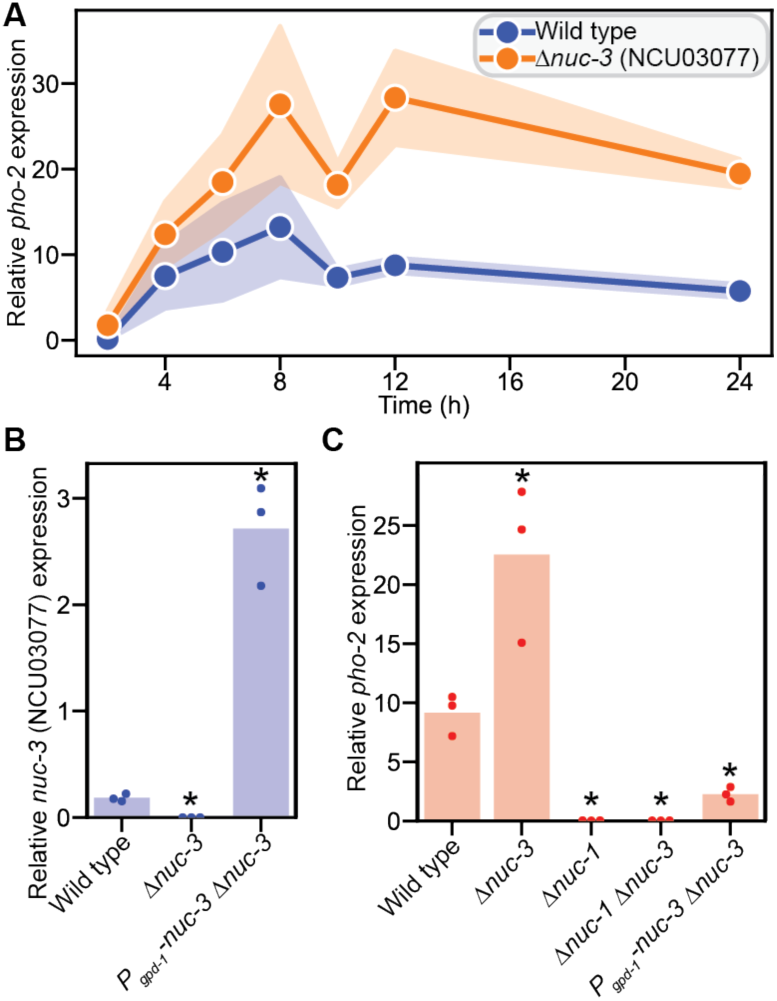
The uncharacterized transcription factor NUC-3 (NCU03077) represses the expression of the alkaline phosphatase *pho-2* during phosphate starvation. **(A)** Time course of *pho-2* expression relative to *act* in the indicated strains. Lines are the mean of at least three biological replicates. Standard deviation is indicated with colored bars. The expression of *pho-2* in the Δ*nuc-3* cells is statistically significantly different than in wild type (*nuc-3^+^*) cells as determined by a 2-way ANOVA (p=5.3×10^-3^). **(B-C)** Expression of *nuc-3* (B) or *pho-2* (C) relative to *act* in the indicated strains. Bars are the mean of three biological replicates (dots). Asterisks indicate expression values that are statistically significantly different from that seen in wild type cells as determined by a Student’s *t*-test with a Benjamini-Hochberg multiple hypothesis correction, *p_adj_<0.05.

The expression of NCU03077 increased in response to phosphate starvation (Fig 1A, 2B, and Dataset S1). Thus, we hypothesized that increasing NCU03077 expression would result in decreased *pho-2* expression. To test this hypothesis, we overexpressed NCU03077 under the control of the *gpd-1* (NCU01528) promoter. The resulting *P_gpd-1_-*NCU03077 ΔNCU03077 strain exhibited NCU03077 expression 15-fold higher than wild type 12h after a shift to phosphate starvation (Fig 3B). Overexpression of NCU03077 resulted in a 4-fold reduction in *pho-2* expression as compared to wild type cells 12h after a shift to phosphate starvation (Fig 3C). These data indicated that NCU03077 repressed expression of *pho-2* after cells experienced phosphate starvation for an extended period. Thus, we named NCU03077, *nuc-3*.

Because *nuc-3* expression was downregulated in Δ*nuc-1* cells relative to *nuc-1^+^* cells and acted later in the phosphate starvation response than NUC-1, we hypothesized that *nuc-1* is epistatic to *nuc-3* (Fig 2B, 3A, and Dataset S1). To test this hypothesis, we deleted *nuc-3* in Δ*nuc-1* cells and measured *pho-2* expression 12h post-shift to phosphate starvation. As expected, Δ*nuc-1* cells showed a 300-fold decrease in *pho-2* expression as compared to wild type cells, while deletion of *nuc-3* resulted in a 2.5-fold increase in *pho-2* expression 12h post-shift to phosphate starvation as compared to wild type (Fig 3C). Expression of *pho-2* in Δ*nuc-1* Δ*nuc-3* cells was indistinguishable from *pho-2* expression in Δ*nuc-1* cells (Fig 3C). These data supported our hypothesis that *nuc-1* was epistatic to *nuc-3*, and NUC-3 repressed *pho-2* either directly or indirectly during phosphate starvation.

### NUC-3 directly repressed genes involved in phosphate acquisition and liberation of phosphate from the cell

The upregulation of *pho-2* in the Δ*nuc-3* mutant relative to wild type (*nuc-3^+^*) cells led us to hypothesize that NUC-3 is responsible for repressing the expression of genes involved in the phosphate starvation response globally. To test this hypothesis, we used RNAseq to measure the transcriptome of *nuc-3^+^* cells and Δ*nuc-3* cells 12h after a shift from 37 mM phosphate to phosphate starvation. Only 24 genes were differentially expressed by at least 4-fold in the Δ*nuc-3* mutant as compared to *nuc-3^+^*cells. Of these 24 genes, 22 were upregulated in Δ*nuc-3* cells relative to *nuc-3^+^*cells (Fig 4A and Dataset S1). Given that *pho-2* expression only increased by 2.5-fold in Δ*nuc-3* cells as compared to wild type during phosphate starvation in our reverse transcription quantitative PCR (RT-qPCR) data (Fig 3A and 3C), we dropped our threshold for differential expression to 2-fold. Ninety genes were at least 2-fold differentially expressed between Δ*nuc-3* cells and *nuc-3^+^*cells 12h after a shift from 37 mM phosphate to phosphate starvation (Fig 4A and Dataset S1). Seventy-nine of these genes were upregulated and 11 genes were downregulated in Δ*nuc-3* cells relative to *nuc-3^+^* cells (Fig 4A and Dataset S1).

**Fig 4.**
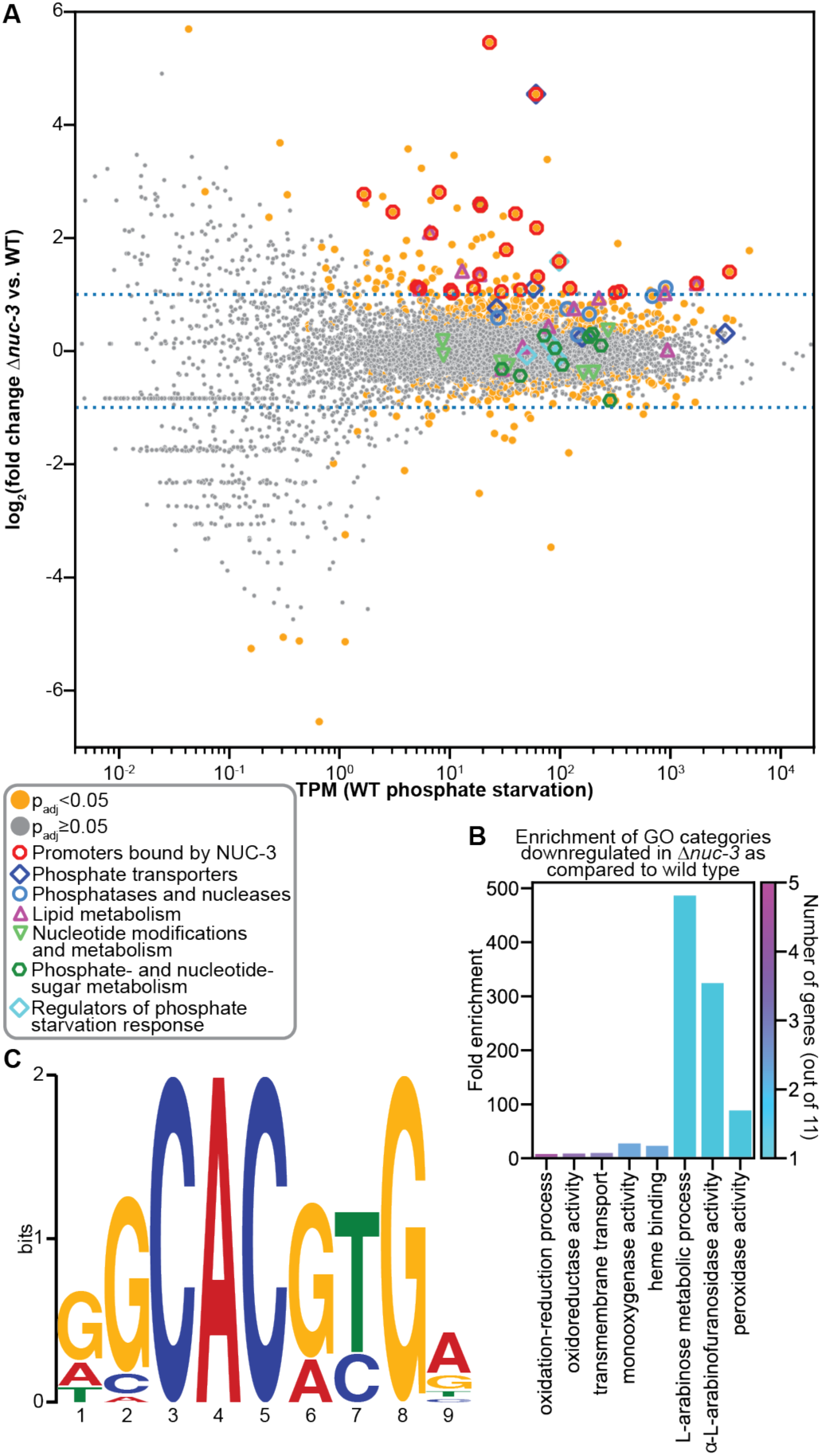
NUC-3 directly represses the expression of phosphate acquisition genes during phosphate starvation. **(A)** Differential expression analysis of Δ*nuc-3* cells as compared to wild type (WT) (*nuc-3^+^*) cells exposed to phosphate starvation. Genes with significant differential expression (p_adj_ < 0.05) are indicated with orange circles. Genes without a significant difference in expression (p_adj_ ≥ 0.05) are indicated in grey. Genes whose promoters are bound by NUC-3 and are differentially expressed by at least 2-fold in the Δ*nuc-3* mutant as compared to wild type (*nuc-3^+^*) cells exposed to phosphate starvation are indicated with red octagons. Genes encoding known or predicted phosphate transporters are indicated with dark blue diamonds. Genes encoding known or predicted phosphatases and nucleases are indicated with light blue circles. Genes predicted to be involved in lipid metabolism are indicated with magenta triangles. Genes predicted to play a role in nucleotide modification or metabolism are indicated with light green triangles. Genes involved in phosphate-sugar and nucleotide-sugar metabolism are indicated with dark green hexagons. Genes encoding regulators of the phosphate starvation response are indicated with light blue diamonds. Dotted blue lines indicated a 2-fold change in expression. The *nuc-3* gene was left off the scatterplot to make it easier to visualize the fold change of the rest of the genes because the large change in *nuc-3* expression was due to the deletion of this gene in the Δ*nuc-3* strain, rather than due to any potential regulation of *nuc-3*. **(B)** Fold enrichment of genes significantly downregulated by at least 2-fold in the Δ*nuc-3* mutant as compared to wild type (*nuc-3^+^*) cells during phosphate starvation in significantly enriched (p_adj_ < 0.05) GO categories. The number of genes significantly downregulated by at least 2-fold found in each category is indicated by the color of the bar. (No GO categories had a significant enrichment in the set of genes upregulated by at least 2-fold in the Δ*nuc-3* mutant as compared to wild type (*nuc-3^+^*) cells during phosphate starvation.) GO enrichment analysis was calculated using FungiFun 2.2.8 (https://elbe.hki-jena.de/fungifun/). **(C)** NUC-3 consensus DNA binding motif (E-value = 1.1 x 10^-5^) of NUC-3 promoter binding sites in genes differentially expressed by at least 2-fold in the Δ*nuc-3* mutant as compared to wild type cells during phosphate starvation built using MEME version 5.5.7 (78).

The 79 genes upregulated in Δ*nuc-3* cells did not show enrichment for any GO categories. However, the 11 genes downregulated in Δ*nuc-3* cells as compared to *nuc-3^+^*cells were enriched for oxidoreductase activity, transmembrane transport, monooxygenase activity, heme binding, arabinose metabolic processes, and peroxidase activity (Fig 4B). Three transporters were downregulated in Δ*nuc-3* cells relative to *nuc-3^+^* cells: the rhamnose transporter *sut-28*, which was also downregulated in Δ*nuc-1* cells relative to *nuc-1^+^* cells; a predicted high-affinity nicotinic acid transporter (NCU09698); and an MFS transporter (NCU06341) (Dataset S1). Given that NUC-1 directly repressed expression of genes associated with the ribosome, we hypothesized that genes activated by NUC-3 would include genes encoding ribosomal proteins or genes associated with ribosome biogenesis. However, none of the genes whose expression was downregulated in Δ*nuc-3* cells by at least 2-fold were predicted to play a role in ribosome biogenesis or transcription.

Most genes regulated by NUC-3 in response to phosphate starvation were upregulated in Δ*nuc-3* cells relative to *nuc-3^+^* cells (Fig 4A). Of the 79 genes upregulated in Δ*nuc-3* cells by at least 2-fold, 21 were activated by at least 4-fold in response to phosphate starvation as compared to 7 mM phosphate in wild type cells and/or were downregulated in Δ*nuc-1* cells as compared to *nuc-1^+^* cells by at least 4-fold during phosphate starvation (Fig 2A). A number of these genes have a clear role in the phosphate starvation response: *nuc-2, pho-2*, *gdp-1*, *bet-6*, NCU09767, and *grn* (Fig 4A and Dataset S1). Several of the genes most highly upregulated in Δ*nuc-3* cells as compared to *nuc-3^+^* cells were also downregulated in Δ*nuc-1* cells as compared to *nuc-1^+^* cells. For example, the MFS phosphate transporter *pho-7* was upregulated by 23-fold in Δ*nuc-3* cells as compared to *nuc-3^+^* cells and downregulated by 4.5-fold in Δ*nuc-1* cells as compared to *nuc-1^+^* cells and a phospholipase domain containing protein (NCU05859) was upregulated by over 4-fold in Δ*nuc-3* cells relative to *nuc-3^+^* cells and in wild type cells exposed to phosphate starvation as compared to 7 mM phosphate (Fig 2A, 4A, and Dataset S1).

Given the role of NUC-3 in repressing the expression of genes in the phosphate starvation response, we hypothesized that NUC-3 is a transcriptional repressor and that genes activated by NUC-3 were due to indirect regulation. To test this hypothesis, we used DAPseq to identify promoter regions bound by NUC-3. We identified 1391 NUC-3 binding sites within 3,000 bp upstream of the translational start site of 1793 genes (Dataset S2). To distinguish between promoter regions bound *in vitro* and genes directly regulated by NUC-3, we compared the genes with NUC-3 promoter binding sites to the genes differentially expressed by at least 2-fold in Δ*nuc-3* cells as compared to *nuc-3^+^* cells after 12h of phosphate starvation. Twenty-six genes fulfilled these criteria (Fig 2A, 4A, and Datasets S1 and S2). In addition, NUC-3 bound its own promoter (Dataset S2). We used these 24 promoter binding sites in these 27 genes to identify GGCACGTGA as the consensus binding motif for NUC-3, which includes the CACGTG motif found in a number of basic helix-loop-helix transcription factor binding motifs, including NUC-1 (52) (Fig 2C and 4C).

All 26 genes that were directly regulated by NUC-3 were repressed in *nuc-3^+^* cells relative to Δ*nuc-3* cells during phosphate starvation, providing additional evidence for the role of NUC-3 as a transcriptional repressor (Fig 2A, 4A, and Dataset S1). These genes included three genes known to play a role in the phosphate starvation response: the alkaline phosphatase *pho-2*, the glycerophosphoryl diester phosphodiesterase *gdp-1*, and the cyclin-dependent kinase inhibitor *nuc-2* (Fig 4A and Dataset S1). NUC-3 also directly repressed several additional genes that may play a role in phosphate acquisition or mobilization, including *pho-7* (Fig 4A and Dataset S1). Cells can liberate phosphate from polyphosphate stores in the vacuole during phosphate starvation (22, 23). Two genes associated with the vacuole were directly repressed by NUC-3. NCU03950 is homologous to vacuolar membrane polyphosphate polymerases that catalyze the synthesis of inorganic polyphosphate (53), and *aap-17* (NCU08066) is homologous to vacuolar amino acid transporters (54). Three additional genes that were directly repressed by NUC-3 may play a role in phosphate scavenging: the UPD-glucose-6-dehydrogenase *gld-2* (NCU08228); *gt2-4* (NCU08226), a gene that is homologous to nucleotide-diphospho-sugar transferases; and NCU06524, which has homology to protease inhibitors and phosphatidylethanolamine-binding proteins (Fig 4A and Dataset S1).

A potential connection to the phosphate starvation response for the remaining 17 genes directly repressed by NUC-3 was less clear. Three transcription factors were directly repressed by NUC-3: the zinc binuclear cluster transcription factor *far-2* (NCU03643) is involved in regulating fatty acid utilization (55, 56); *tcf-10* (NCU05994), a transcription factor homologous to Dal81/TamA, which is involved in regulating nitrogen metabolism in *A. nidulans* (57, 58); and NCU04630 encodes a putative C2H2-type domain containing protein. Aside from *pho-7* and *aap-17*, two additional transporters were directly repressed by NUC-3: the predicted quinate transporter *mfs-12* (NCU05585) and the calcium-transporting ATPase *trm-1* (NCU07966). NUC-3 directly repressed two genes with potential roles in regulating conidiation: *con-6* (NCU08769) (59) and the integral membrane protein NCU00848 (60). NUC-3 also repressed the anchored cell wall protein *acw-6* (NCU03530), the malate/L-lactate dehydrogenase NCU05586, the F-box domain containing protein NCU08842, and the predicted molybdenum cofactor sulfurase *nit-14* (NCU03011). The remaining six genes directly repressed by NUC-3 encoded hypothetical proteins: NCU02722, NCU03078, NCU07485, NCU07967, NCU08053, and NCU08282 (Fig 4A and Dataset S1).

## DISCUSSION

Phosphate is a key component of nucleic acids, phospholipids, and other cellular metabolites, making it critical for cellular function (1). In the soil, carbon is abundant, but phosphate and other nutrients are limiting (61, 62). Thus, fungal cells have evolved mechanisms to increase phosphate acquisition and liberate phosphate from cellular molecules when extracellular phosphate levels drop (1). The response to phosphate limitation is broadly conserved amongst ascomycete fungi (9) and is activated by the basic helix-loop-helix transcription factor NUC-1 (27). Previously, microarrays were used to investigate transcriptional changes in wild type *N. crassa* cells exposed to phosphate limitation (32). We used RNAseq and DAPseq to examine the global phosphate starvation response in *N. crassa* and the role of NUC-1. Approximately two thirds of the genes identified as at least 4-fold differentially expressed in response to phosphate limitation via microarray were also differentially expressed in our RNAseq dataset (Dataset S1) (32).

NUC-1 was identified through classical genetics for its role in activating the phosphate starvation response (27). Our RNAseq and DAPseq data showed that of the 27 NUC-1 direct targets, approximately half are activated and half repressed by NUC-1 (Fig 5). This data suggests that NUC-1 acts as a bifunctional transcription factor to directly regulate both the activation of genes involved in phosphate acquisition or liberation and the repression of genes involved in phosphate-hungry cellular processes, such as ribosome biogenesis and transcription. Although examples of bifunctional transcription factors in fungi are limited, *N. crassa* CRE-1 acts as an activator during carbon starvation and a repressor when preferred carbon sources are present (40, 63), and *Aspergillus fumigatus* conidial melanin production is regulated by bifunctional transcription factors (64).

**Fig 5.**
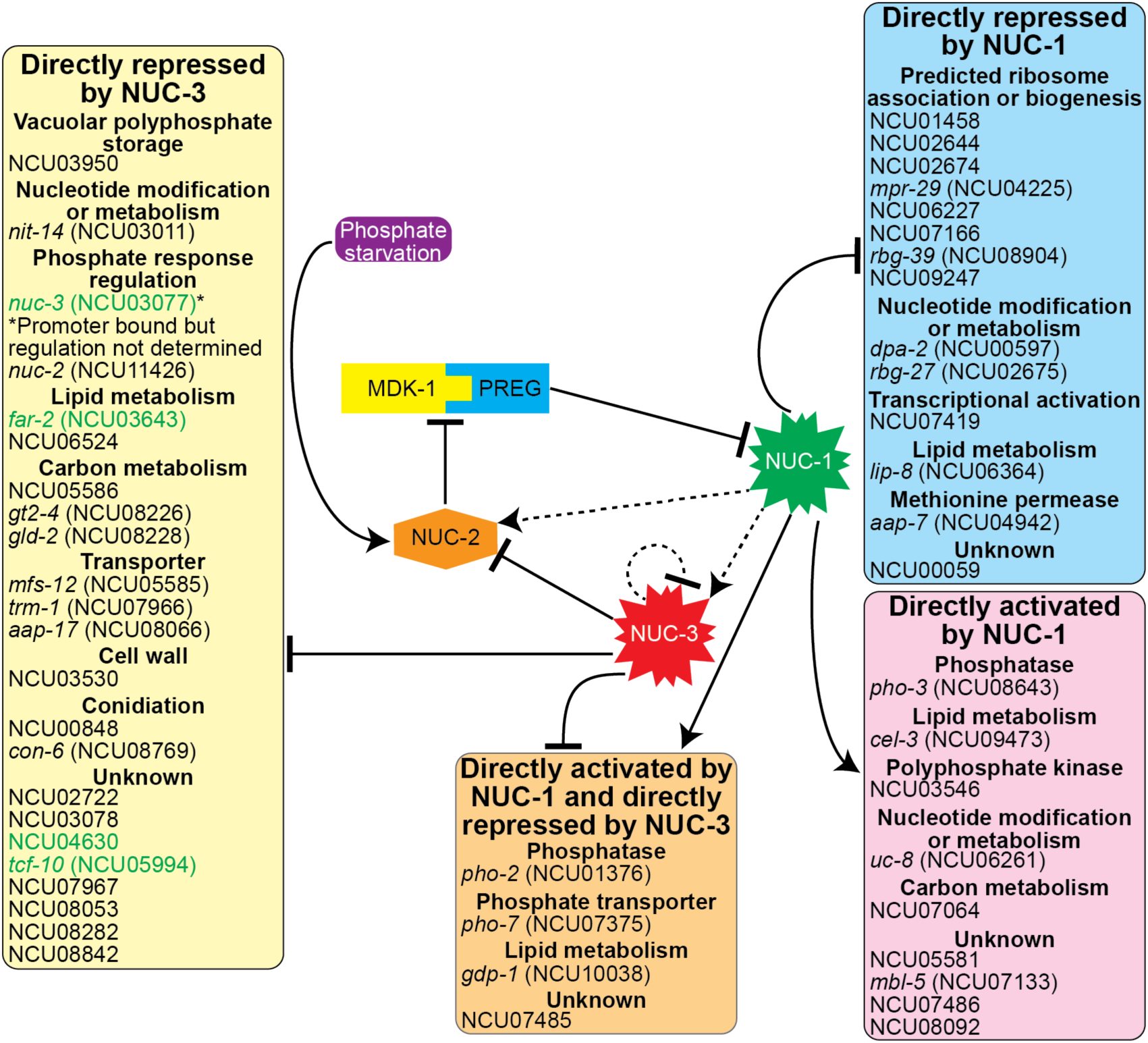
Model for the role of NUC-3 as a brake in the phosphate starvation response regulatory network. When phosphate is abundant a complex comprised of the cyclin-dependent kinase MDK-1 and the cyclin PREG represses the activity of the transcription factor NUC-1 by restricting NUC-1 localization to the cytoplasm. When phosphate is limiting, NUC-2 represses the activity of the MDK-1/PREG complex and NUC-1 translocates to the nucleus where it directly activates the expression of 13 genes and directly represses the expression of 14 genes. NUC-1 indirectly activates the expression of *nuc-2*, which may limit cytoplasmic localization of NUC-1 during phosphate limitation. NUC-1 also indirectly activates the expression of the transcription factor *nuc-3*. When NUC-3 is expressed, it directly represses the expression of 26 genes. NUC-3 also binds its own promoter, but self-regulation of *nuc-3* has yet to be definitively determined. These repressed genes include a number involved in the phosphate starvation response, including *pho-2*, *pho-7*, *gpd-1*, NCU03950, and *nuc-2*. Repression of NUC-2 will, in turn, release the inhibition of the MDK-1/PREG complex, causing increased inhibition of NUC-1 activity and act as a brake during the phosphate starvation response. While gene regulation by NUC-1 occurs relatively quickly after phosphate starvation begins, phosphate-responsive gene regulation by NUC-3 does not occur until cells have experienced phosphate starvation for at least 10h. This may limit the phosphate-intensive activities of transcription and translation when phosphate starvation persists for extensive periods of time. NUC-3 may also act to indirectly repress a number of other processes through the direct repression of 3 additional genes with DNA binding domains (indicated with green text). Solid lines indicate direct regulation (either direct protein interactions or direct binding to promoter regions). Dotted lines indicate regulation for which no direct interaction is known.

The number of genes differentially expressed in response to phosphate starvation is much larger than the number of genes directly regulated by NUC-1 (Fig 2A and Dataset S1). It is possible that NUC-1 binds the promoters of additional genes *in vivo* where chromatin state and the presence of other transcriptional activators and repressors could affect NUC-1 promoter binding. It is also possible that additional transcriptional regulators of the phosphate starvation response exist that have yet to be identified. While *nuc-3* was the only transcription factor whose expression changed by at least 4-fold in the Δ*nuc-1* mutant as compared to wild type cells, our DAPseq data identified NUC-1 binding sites in the promoters of two additional transcription factors that were not regulated by NUC-1 in our RNAseq data: *tcf-3* (NCU00223), homologous to *meaB*, a regulator of nitrogen catabolite repression in *A. nidulans* (65), and *tcf-24* (NCU03273) (Dataset S2). Aside from *nuc-3*, there were seven additional transcription factors that were activated in response to phosphate starvation in wild type cells: NCU01074, NCU06965, NCU05909, *sah-1* (NCU04179), *tah-1* (NCU00282), *xlr-1* (NCU06971), and *znf-10* (NCU05767) (Dataset S1). The roles of most of these transcription factors are uncharacterized, but *xlr-1* is critical for activating genes necessary to utilize hemicellulose (66, 67). These data suggest that the phosphate starvation response may be linked to responses to other nutrient classes as has been seen in the case of responses to carbon, nitrogen, and sulfur (40, 44, 45, 67-69). It will be the work of future studies to identify any potential roles for these transcription factors in the phosphate starvation response.

While transcriptional activation of phosphate acquisition and liberation genes is necessary to improve the chances of cellular survival during phosphate starvation, transcription followed by translation requires phosphate. NUC-1 directly represses genes associated with the ribosome, ribosome biogenesis, and transcriptional activation (Fig 5). Additionally, the presence of NUC-3 in the phosphate starvation regulatory circuit may explain how cells limit the phosphate utilized by transcription when cells must survive extended phosphate starvation. NUC-3 is conserved among filamentous ascomycete fungi, suggesting this model for regulating the phosphate starvation response could exist in many fungi (Fig S2).

We demonstrated that the transcription factor *nuc-3* is activated in response to phosphate starvation in a NUC-1-dependent fashion (Fig 1A, 2A, 2B, 5, and Dataset S1). When NUC-3 is expressed, it represses expression of genes associated with phosphate acquisition, transport, and storage and liberation of phosphate from cellular stores (Fig 5). NUC-3 also represses expression of the cyclin-dependent kinase inhibitor *nuc-2* (Fig 5). Reduced levels of NUC-2 could result in released inhibition of the MDK-1/PREG complex (28, 33). This complex would then be free to inhibit the activity of NUC-1 and further reduce expression of NUC-1 target genes (Fig 5). Thus, NUC-3 acts as a brake on both expression of specific genes in the phosphate starvation response and on the activity of NUC-1, the main activating transcription factor of the phosphate starvation response, when phosphate starvation persists even after the phosphate starvation response is fully deployed (Fig 5). This regulatory circuit presents a model for how the phosphate transcriptional network balances the cellular requirement to increase phosphate availability while limiting transcription and translation to save phosphate for other essential cellular processes.

## MATERIALS & METHODS

### N. crassa strains and culturing

Strains used in this study are listed in Table S1. All strains were derived from the wild type reference strains FGSC 2489 (*mat A*) and FGSC 4200 (*mat a*) using standard genetic techniques and confirmed by PCR and DNA sequencing (70, 71). *N. crassa* cells were grown from freezer stocks on Vogel’s minimal medium + 2% sucrose + 1.5% agar (Thermo Fisher Scientific) slants (72) for 2 d at 30°C in the dark and 4 to 8 d at 25°C in constant light. *N. crassa* conidia were harvested from slants with sterile double distilled H_2_O for inoculation. All chemicals were purchased from Sigma-Aldrich unless otherwise noted.

### RNA sequencing and transcript abundance

The indicated strains were inoculated into 3 mL Vogel’s minimal medium (72) in round bottom, deep well 24-well plates at 10^6^ conidia/mL and grown for 24 h at 25°C in constant light with 200 rpm shaking. The media was then vacuumed out of the wells and mycelial mats were washed three times with either 3 mL Fries minimal medium (73, 74) containing 7 mM phosphate or lacking phosphate and then shifted to 3 mL Fries minimal medium containing 7 mM phosphate or lacking phosphate, respectively. Cells were then incubated for the indicated amount of time at 25°C in constant light with 200 rpm shaking.

Mycelia were harvested by filtering on Whatman paper no. 1 and flash frozen in liquid nitrogen for RNAseq and RT-qPCR. RNA extraction and RNAseq library preparation were performed as described in Wu *et al* (2020) (40). Sequencing conditions are listed in Table S2. For details see SI Materials and Methods.

The transcript abundance for RNAseq (transcripts per million, TPM) was quantified using Salmon v. 1.4.0 (75) mapping to the *N. crassa* OR74A genome (v12) (76). Differential expression was determined using DESeq2 v. 1.44.0 (77). For details see SI Materials and Methods. RT-qPCR was performed to determine relative transcript abundance with *act* (NCU04173) as the housekeeping control gene using the EXPRESS One-Step SYBR GreenER kit (Life Technologies). RT-qPCR primer sequences are in Table S3.

### Statistical significance tests

All experiments had at least 3 biological replicates, which refers to independently inoculated cultures. Statistical significance was determined using DESeq2 v1.44.0 for RNAseq experiments (77). Statistical significance for the RT-qPCR experiments was determined using a 2-way ANOVA test for the time course (Fig 3A) and a two-tailed homoscedastic (equal variance) Student’s *t*-test with a Benjamini-Hochberg multiple hypothesis correction for all other experiments. In bar graphs, bars indicate the mean of biological replicates and dots indicate the individual biological replicates.

### DAPseq

DAPseq was performed as described in (40). For details see SI Materials and Methods.

### DNA binding consensus motif generation

Motif discovery was performed using Multiple Expectation maximizations for Motif Elicitation (MEME) v5.5.5 (NUC-1) or v5.5.7 (NUC-3) (78). For details see SI Materials and Methods.

### Functional enrichment analysis

Functional enrichment analysis was done using the FungiFun2 online resource tool (https://elbe.hki-jena.de/fungifun/). For details see SI Materials and Methods.

### NUC-3 phylogenetic tree generation

The Basic Local Alignment Search Tool for proteins (BLASTP) was used to identify NUC-3 homologs. Identified homologs were aligned with MAFFT v7.487 (79). The phylogenetic tree was constructed using FastTree v2.1.8 (80). For details see SI Materials and Methods.

### Data availability

RNAseq data used in this study were deposited in the Gene Expression Omnibus (GEO) at the National Center for Biotechnology Information (NCBI) and are accessible through GEO series accession number GSE293601. Processed RNAseq data are available in Dataset S1. DAPseq data used in this study were deposited in the NCBI Sequence Read Archive (SRA) and are accessible through SRA series accession number PRJNA436200. Processed DAPseq data are available in Dataset S2. The numerical values used to generate all RT-qPCR graphs are shown in Dataset S3. Strains constructed in this study are available from the Fungal Genetics Stock Center (www.fgsc.net) (71).

## Supporting information

Supplemental Appendex

## ACKNOWLEDGEMENTS

This work was supported by an Energy Biosciences Institute grant, a Laboratory Directed Research and Development Program of Lawrence Berkeley National Laboratory grant under US Department of Energy (DOE) Contract DE-AC02-05CH11231, a Joint Genome Institute (JGI) Community Science Program grant (CSP 982), an Innovative Genomics Institute grant, and funds from the Fred E. Dickinson Chair of Wood Science and Technology to N.L.G. This work was supported by a grant from the National Institute of General Medical Sciences of the National Institutes of Health (NIH) under Award Number R35GM150926 to L.B.H. V.W.W. was partially supported by NIH National Research Service Award Trainee Grant 5T32GM007127-39. The work (proposal 10.46936/10.25585/60001007) conducted by the DOE JGI (https://ror.org/04xm1d337), a DOE Office of Science User Facility, was supported by the Office of Science of the US DOE under Contract DE-AC02-05CH11231.

