## Supplemental Appendex for "A novel regulator of the fungal phosphate starvation response revealed by transcriptional profiling and DNA affinity purification sequencing"

**SI MATERIALS and METHODS**

*RNA sequencing and transcript abundance*

The indicated strains were inoculated into 3mL Vogel's minimal medium (1) in 24-well plates at $10^6$ conidia/mL and grown for 24 h at 25°C in constant light with constant shaking at 200 rpm. The media was then vacuumed out of the wells and mycelial mats were washed three times with either 3 mL Fries minimal medium (2, 3) containing 7 mM phosphate or Fries minimal medium lacking phosphate and then shifted to 3 mL of either Fries minimal medium containing 7 mM phosphate or Fries minimal medium lacking phosphate, respectively. Cells were then incubated for the indicated amount of time at 25°C in constant light with constant shaking at 200 rpm.

Mycelia were harvested by filtering on Whatman paper no. 1 and flash frozen in liquid nitrogen. RNA extraction and library preparation were performed as described in Wu *et al* (2020) (4). Briefly, total RNA was harvested from mycelia flash frozen in liquid nitrogen the indicated amount of time post-transfer to the indicated media using a TRIzol (Life Technologies) extraction and cleaned up with the RNeasy kit (Qiagen). Libraries were prepared from total RNA using poly(A) enrichment and standard Illumina protocols. Libraries were prepared and sequenced at the Joint Genome Institute on the Illumina HiSeq platform with 100 bp single end reads or at the University of California Berkeley Vincent J. Coates Genomics Sequencing Lab on the Illumina HiSeq platform with 50bp single end reads or NovaSeq platform with 150 bp paired end reads as listed in Table S2.

The transcript abundance (transcripts per million, TPM) was quantified using Salmon v. 1.4.0 mapping to the *N. crassa* OR74A genome (v12) (5) with the --validateMappings setting (6). Differential expression was determined using DESeq2 v. 1.44.0 (7). Differential expression was only called between samples sequenced at the same center and sequencing platform. Genes were denoted as differentially expressed between wild type exposed to Fries minimal medium with 7 mM phosphate as compared to Fries minimal medium lacking phosphate and wild type (*nuc-1*⁺) cells as compared to Δ*nuc-1* cells exposed to Fries minimal medium lacking phosphate if there was at least a 4-fold change in expression and the TPM of the gene was at least 10 in either of the 2 conditions. Genes were denoted as differentially expressed between wild type (*nuc-3*⁺) cells as compared to Δ*nuc-3* cells exposed to Fries minimal medium lacking phosphate if there was at least a 2-fold change in expression and the TPM of the gene was at least 10 in either of the 2 conditions.

RNAseq data used in this study were deposited in the Gene Expression Omnibus (GEO) at the National Center for Biotechnology Information (NCBI) and are accessible through GEO series accession number GSE293601. Processed RNAseq data are available in Dataset S1.

*DAPseq*

DAPseq was performed as described in (4). Briefly, cDNA was generated from RNA harvested from the wild type FGSC 2489 using the EcoDry premix (Clontech). Predicted open reading frames for *nuc-1* (NCU09315) and *nuc-3* (NCU03077) were amplified and inserted into an expression vector upstream of a HALO tag. NUC-1 and NUC-3 proteins were then produced using the Promega TnT T7 Rabbit Reticulocyte Quick Coupled Transcription/Translation System by incubating 1 μg of plasmid DNA with 60 μL of TnT Master Mix and 1.5 μL of 1 mM methionine overnight at room temperature. NUC-1 and NUC-3 protein expression was verified by Western blot using Promega anti-HaloTag monoclonal antibody.

The NUC-1 and NUC-3 transcription and translation reactions were incubated with 1 μg salmon sperm for blocking, 20 μl Promega Magne HaloTag Beads, and 20 ng of genomic DNA libraries prepared with the KAPA library kit for Illumina sequencing from genomic DNA harvested from the wild type FGSC 2489 strain grown on liquid Vogel's minimal medium for 24 h at 25°C, sheared to a 300 bp peak using a Covaris LE220 sonicator, and size selected using AMPure XP beads on a rotator for 1 h at room temperature. The bead-bound protein and protein-bound DNA were then washed three times with 2.5% Tween20 in phosphate buffered saline, resuspended in 30 μL ddH2O, and heated to 98°C for 10 m to denature proteins and release bound DNA into solution. The supernatant containing any released DNA was transferred to a new tube for PCR amplification using KAPA Hifi polymerase for 12-16 cycles to generate DAPseq DNA libraries. A DAPseq DNA library was generated in the same conditions with no plasmid added to the TnT Master Mix as a negative control. Single DAPseq libraries were generated for NUC-1 and NUC-3 and sequenced with 150 bp paired end reads on an Illumina MiSeq.

Filtered reads were aligned to the *N. crassa* OR74A genome (v12) using Bowtie v2.3.2 (8). Peak calling was performed using MACS v2.1.1 with p-value cutoff at 0.001 and utilizing negative control library alignments (9). Peaks within 3000 bp upstream of translation start sites were selected for and annotated with a custom Python script. DAPseq data was deposited in the NCBI Sequence Read Archive (accession number PRJNA436200; ID SRP133627).

*DNA binding consensus motif generation*

Motif discovery was performed using Multiple Expectation maximizations for Motif Elicitation (MEME) v5.5.5 (NUC-1) or v5.5.7 (NUC-3) (10). The input for MEME motif discovery was DAPseq binding peak sequences with a maximum motif width of 50 bp, a minimum motif width of 5 bp, any number of motif sites in the sequences, the classic objective function, and a $0^{th}$ order Markov model for sequences. Binding peak sequences were included in motif generation for NUC-1 or NUC-3 if they were within 3000 bp upstream of a translational start site that was at least 2-fold differentially expressed between wild type and either the Δ*nuc-1* or Δ*nuc-3* mutant, respectively, during exposure to Fries minimal medium lacking phosphate and had a TPM of at least 10 in either wild type or the Δ*nuc-1* or Δ*nuc-3* mutant, respectively.

*Functional enrichment analysis and gene annotation*

Functional enrichment analysis was done using the FungiFun2 online resource tool (https://elbe.hki-jena.de/fungifun/) with KEGG as the classification ontology (11, 12). The gene to category associations were tested for overrepresentation using hypergeometric distribution with Benjamini-Hochberg correction for false discovery rate.

Gene annotations were pulled from FungiDB (https://fungidb.org) (13) or inferred from homology to characterized genes in related fungi.

*NUC-3 phylogenetic tree generation*

The Basic Local Alignment Search Tool for proteins (BLASTP) was used to identify NUC-3 homologs using the reference proteins database of the indicated species and the following settings: blastp algorithm, 100 max target sequences, short queries, 0.05 expect threshold, a word size of 4, 0 max matches in query range, the BLOSUM62 matrix, gap costs of existence 11 and extension 1, the conditional compositional score matrix adjustment, and filtering low complexity regions.

Identified homologs were aligned with MAFFT v7.487 using the FFT-NS-2 strategy (14). The phylogenetic tree was constructed using FastTree v2.1.8 using the maximum likelihood method based on the Jones-Taylor-Thornton matrix-based model with *Candida albicans* Try6p as the outgroup (15). The reliability of each split in the tree was computed using the Shimodaira-Hasegawa test on three alternate topologies around that split.

### SI FIGURES

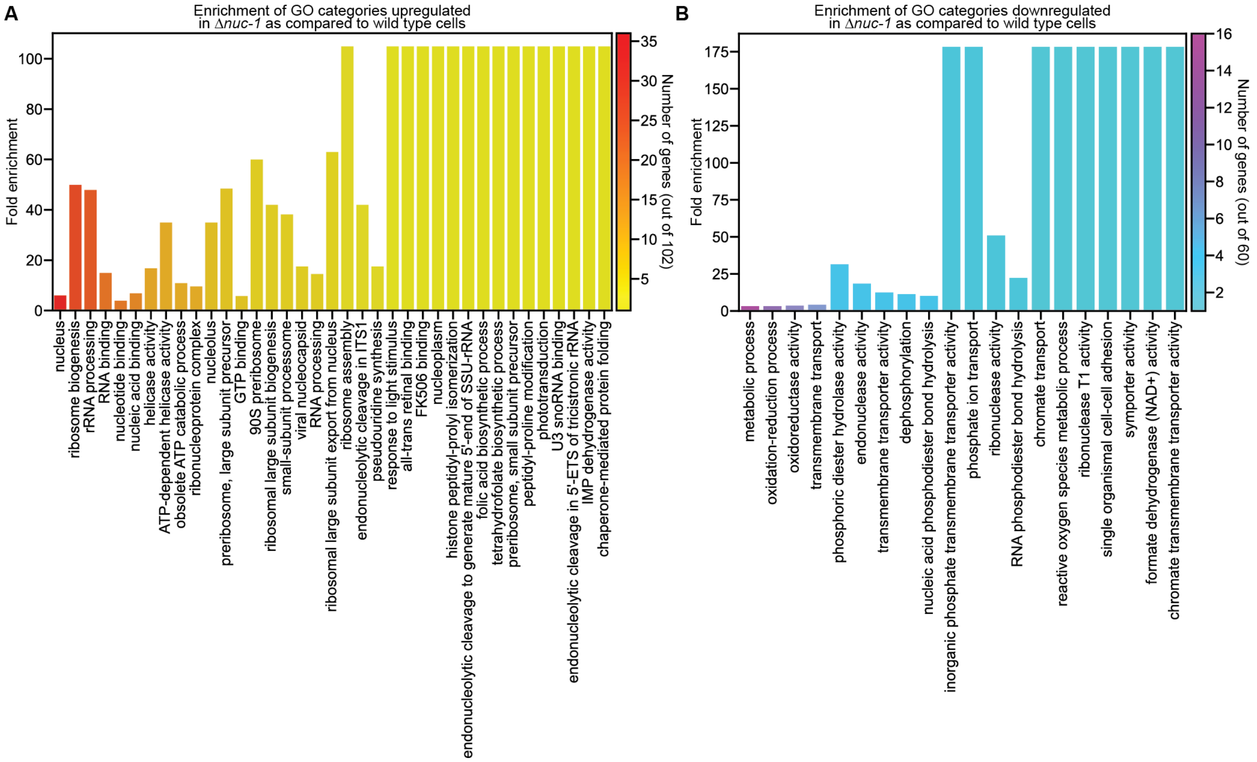


**Fig S1. The transcription factor NUC-1 activates genes involved in phosphate acquisition and liberation and represses genes associated with the ribosome and transcription. (A)** Fold enrichment of genes significantly upregulated by at least 4-fold in the Δ*nuc-1* mutant as compared to wild type (*nuc-1*$^{+}$) cells during phosphate starvation in significantly enriched ($p_{adj}$ < 0.05) gene ontology (GO) categories. The number of genes significantly upregulated by at least 4-fold found in each category is indicated by the color of the bar. **(B)** Fold enrichment of genes significantly downregulated by at least 4-fold in the Δ*nuc-1* mutant as compared to wild type (*nuc-1*$^{+}$) cells during phosphate starvation in significantly enriched ($p_{adj}$ < 0.05) gene GO categories. The number of genes significantly downregulated by at least 4-fold found in each category is indicated by the color of the bar. GO enrichment analysis was calculated using FungiFun 2.2.8 (https://elbe.hki-jena.de/fungifun/).

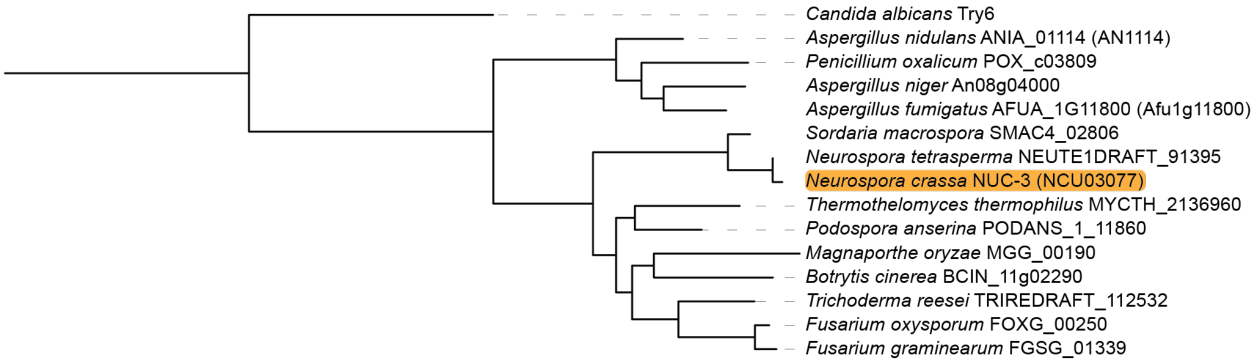


**Fig S2. NUC-3 is conserved in the Pezizomycotina.** Protein sequences of NUC-3 homologs were used to build a phylogenetic tree using the maximum likelihood method based on the Jones-Taylor-Thornton matrix-based model using FastTree with *Candida albicans* as an outgroup (15). The reliability of each split in the tree was computed using the Shimodaira-Hasegawa test on three alternate topologies around that split. *N. crassa* NUC-3 is highlighted in orange.

### SI TABLES

**Table S1. Strains used in this study.**

| Strain name | Genotype | Source |
|---|---|---|
| Wild type | Wild type *mat A* | FGSC* 2489 (16) |
| Wild type | Wild type *mat a* | FGSC 4200 (16) |
| Δ*nuc-1* | Δ*nuc-1* (NCU09315)::*Hyg*$^R$ *mat a* | FGSC 11448 (17) |
| Δ*nuc-3* | Δ*nuc-3* (NCU03077)::*Hyg*$^R$ *mat a* | FGSC 11356 (17) |
| Δ*nuc-3* | Δ*nuc-3*::*Nat*$^R$ *mat A* | This study (FGSC 26801) |
| Δ*nuc-1* Δ*nuc-3* | Δ*nuc-1*::*Hyg*$^R$ Δ*nuc-3*::*Nat*$^R$ *mat a* | This study (FGSC 26800) |
| $P_{gpd-1}$-*nuc-3* Δ*nuc-3* | Δ*csr-1*::$P_{gpd-1}$-*nuc-3* Δ*nuc-3*::*Nat*$^R$ *mat A* | This study (FGSC 26802) |

*FGSC stands for Fungal Genetics Stock Center (16).

**Table S2. RNAseq library construction and sequencing.**

| Strain | Condition | Location | Sequencer | Read Length |
|---|---|---|---|---|
| Wild type *mat A* | 7.3 mM phosphate (4 h) | Joint Genome Institute | HiSeq 2000 | 1x100 bp |
| Wild type *mat A* | 0 mM phosphate (4 h) | Joint Genome Institute | HiSeq 2000 | 1x100 bp |
| Wild type *mat A* | 0 mM phosphate (4 h) | UC Berkeley | HiSeq 4000 | 1x50 bp |
| Δ*nuc-1::HygR mat a* | 0 mM phosphate (4 h) | UC Berkeley | HiSeq 4000 | 1x50 bp |
| Δ*nuc-3::HygR mat a* | 0 mM phosphate (12 h) | UC Berkeley | NovaSeq | 2x150 bp |
| Wild type *mat a* | 0 mM phosphate (12 h) | UC Berkeley | NovaSeq | 2x150 bp |

**Table S3. RT-qPCR primer sequences.**

| Gene | Forward primer 5' → 3' | Reverse primer 5' → 3' |
|---|---|---|
| *act* | TGATCTTACCGACTACCT | CAGAGCTTCTCCTTGATG |
| *nuc-3* | CGGGCCTAACACTGACTAGC | CCTATCCGGTTGCGTCTGTT |
| *pho-2* | TTTACGTTGACTCGTCGCCA | AGCGCCATATTCATACCGGG |

### SI DATASETS

**Dataset S1. TPM counts and differential expression analysis of wild type, Δ*nuc-1*, and Δ*nuc-3* cells exposed to the indicated concentration of phosphate.**

**Dataset S2. DAPseq data for NUC-1 and NUC-3.**

**Dataset S3. RT-qPCR data from wild type and mutant cells exposed to phosphate starvation.**

*Neurospora* reveals functions for multiple transcription factors. Proc Natl Acad Sci U S A 103:10352-10357.
